# Study of the binding mechanisms between palytoxin and its aptamer by docking and molecular simulation

**DOI:** 10.1101/507087

**Authors:** Bo Hu, Rong Zhou, Zhengang Li, Jiaxiang Qin, Shengqun Ouyang, Zhen Li, Wei Hu, Lianghua Wang, Binghua Jiao

**Affiliations:** Department of Biochemistry and Molecular Biology, College of Basic Medical, Second Military Medical University, Shanghai 200433, China; Marine Biological Institute, College of Marine Military Medicine, Second Military Medical University, Shanghai 200433, China; Chengdu FenDi Technology Co., Ltd, Chengdu 610041, China

**Author notes:** Corresponding author, (LW); (BJ). These authors contributed equally to this work (Bo Hu, Rong Zhou, Zhengang Li).

## Abstract

This paper provides a feasible model for aptamer and its target in molecular structure analysis and interaction mechanism. In this study, modeling and dynamic simulation of ssDNA aptamer (P-18S2) and target (Palytoxin, PTX) were performed separately. Then, the combination mechanism of DNA and PTX were predicted, and docking results showed that PTX can combine steadily in the groove at the top of DNA model trough strong hydrogen-bonds and electrostatic interaction. Therefore, we have further truncated and optimized to P-18S2 by simulating, at the same time, we also confirmed the reliability of simulative results by experimenting. Combining with the experimental and computational results, we provide a more reasonable interpretation for the high affinity and specific binding of P-18S2 and PTX and offer the basis for aptamer development in molecular diagnostics and therapeutic application.

**Author summary:** In order to further study the complex structure and interaction of P-18S2 and PTX, a series of molecular modeling program were designed, including modeling, traditional dynamics simulation and molecular docking. Modeling results reveal that the structure of P-18S2 is a DNA G-quadruplex. Meanwhile, the 3D structure of PTX with lowest total energy after equilibrium was selected to use for the subsequent simulations. Then, based on the DNA with G-quadruplex structure, the combination model of DNA and PTX were predicted, and docking results showed that PTX can combine steadily in the groove at the top of DNA model trough strong hydrogen-bonds and electrostatic interaction. Futhermore, we compared the affinity of 6 optimized aptamers by computer simulating to primary P-18S2 bind to PTX respectively, the results showed no significant difference. Therefore, we have further truncated and optimized to P-18S2. In addition, this paper further refined research method based on our previous study^[17]^, the instability of the target structure was considered and optimized, and the biological experiments were used to confirm the veracity of the simulative results. Combining with the experimental and computational methods, we obtained a reasonable interpretation for the high affinity and specific binding of P-18S2 and PTX. In summary, we established a feasible model for aptamer and its target in molecular structure analysis and interaction mechanism, and offer the basis for the study of aptamer development in molecular diagnostics and therapeutic application.

## Introduction

Palytoxin (PTX) is one of the most powerful non-protein natural toxin, which was first separated from soft corals in 1971^[1]^. The LD50 of PTX after intraperitoneal injection is 25 ng/kg and 50 ng/kg in rabbits and mice respectively^®^. Furthermore, PTX can cause dizziness, weakness, muscle pain, breathing difficulties, heart failure, and even death in humans by ingesting seafood contaminated with PTX or direct contact with aerosolized water during dinoflagellate blooms^[3–5]^. Fortunately, the problem of affecting human health and global shellfish industry development due to the contamination of shellfish by PTX has attracted great attention. Meanwhile, many detection methods for PTX have been developed such as mouse bioassay^[6]^, liquid chromatography coupled to a fluorometer, ultraviolet-visible spectrophotometer, or mass spectrometer^[7–9]^. However, there are still many challenges such as the ethical issues and the expensive instruments. We have also been working on PTX research and successfully obtained a DNA aptamer named P18-S2(GGTGGGTCGGACGGGGGTGG), that can bind to PTX with high affinity and specificity, which could serve as a molecular recognition element in diagnosis and biological activity inhibition assays for PTX^[10]^.

Aptamers are functional single-stranded DNA or RNA oligo nucleotides, which are selected from a random oligonucleotide libraries through systematic evolution of ligands by the exponential enrichment (SELEX) technique^[11]^. These aptamers have been isolated and adopted as diagnostic or therapeutic tools, can bind to various targets with high affinity and specificity by folding into steady and particular three-dimensional structures through intermolecular interactions such as the stacking of aromatic rings, electrostatic, van der Waals interactions, hydrogen bonds and induced fit mechanisms^[12, 13]^.The development of aptamers offer a new opportunity to overcome the challenges of traditional methods for detecting toxins and address the risks of seafood and water contaminated with toxins. There are numerous aptamers have been developed into a novel detection method for toxins^[14–16]^. At the same time, the interaction mechanism of aptamers with targets still needs further exploration for the therapeutic application of aptamers. In our previous work, we obtained the aptamer(GO18-T-d)^[15]^ of GTX1/4 and analysed the binding mechanism of GTX1/4 and GO18-T-d by a series of molecular modeling programs^[17]^. In addition, we also obtained the aptamer P18-S2 that can bind to PTX with a high Kd of 0.93 nM^[10]^by SELEX and Biolayer interferometry (BLI) which is a real-time optical analytical technique for measuring interactions between biomolecule^[18]^. In this study, we also designed a series of molecular modeling programs in order to research and further interpret the binding mechanism between P-18S2 and PTX for the therapeutic application of aptamers.

## Results

### Analysis of docking

After modeling and optimization, the 3D structure of DNA P-18S2 with G-quadruplex structure was used as receptor in the docking. The combination of receptor with ligands was evaluated by the Etotal, Eshape and Eforce in the docking results. The electrostatic energy, Eforce, and the steric complementarity score, Eshape, were combined to give a total energy (Etotal) for the complex (in kJ/mol units)^[19]^. In our Hex interaction simulation, the lower value of Etotal would result in more stable combination between DNA and PTX. As is shown in the S1 table, the total calculated interaction energy was listed, which showed the best energy ranked results. The observed best interaction energy and steric complementarity score of binding of DNA P-18S2 with PTX were calculated to be −509.9 kcal mol-1 and −453.4 kcal mol-1 respectively, indicated that the combination of P-18S2 and PTX was extremely stable, and well match in shape (Fig 1). Furthermore, electrostatic energy from P-18S2 and PTX was much lower, led to the higher value of interaction energy and more stable combination between P-18S2 and PTX.

**Figure 1.**
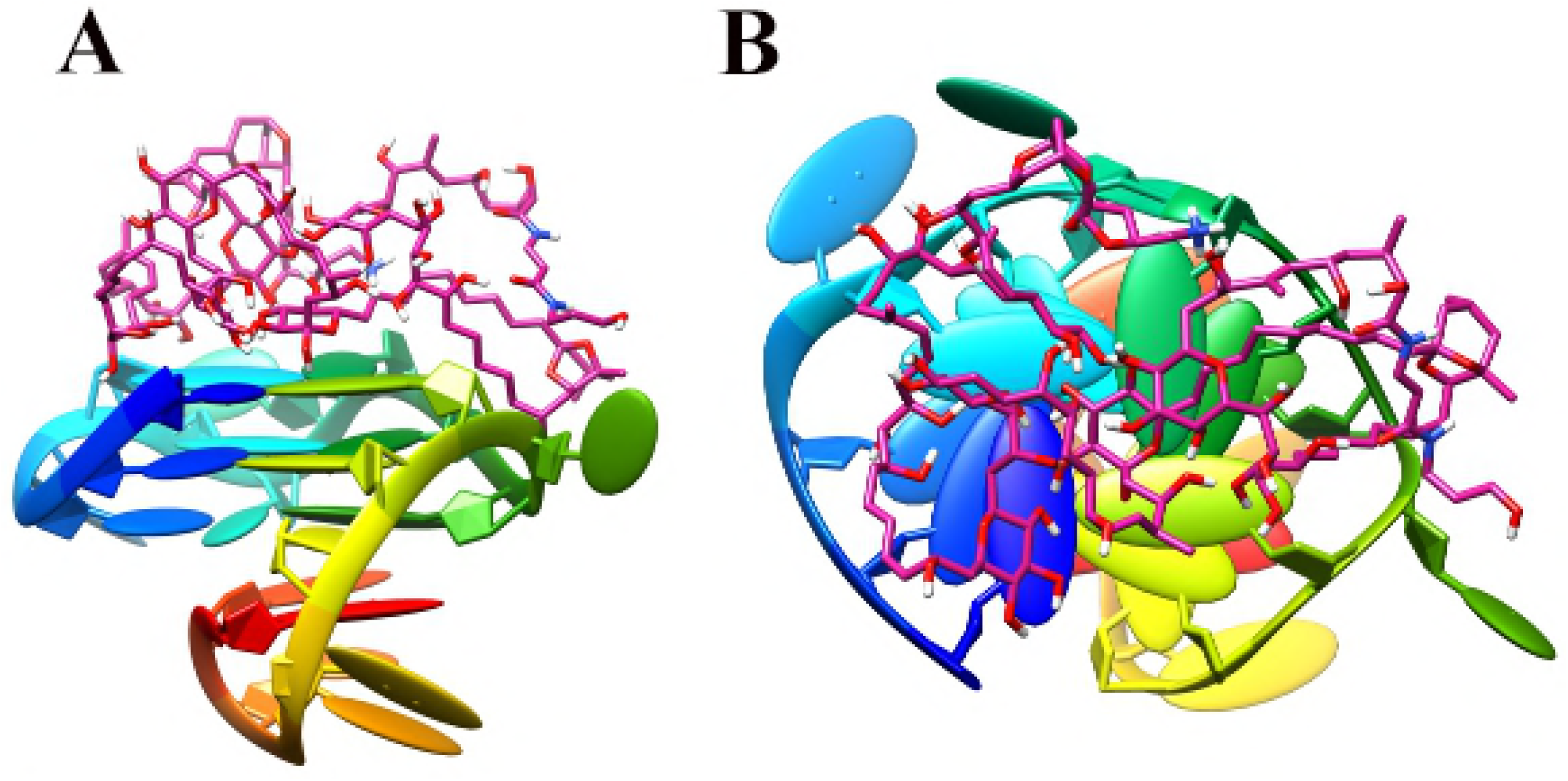
Three dimensional side view (A) and top view (B) of the top-ranking docked conformations between P-18S2 and PTX.

In order to understand the interaction between DNA and ligands, some deeply analysis were performed to the top-ranking docked conformation of complex. As is shown in the Fig 1, the PTX inserted in docking site at the top of DNA P-18S2, made the combination stable. Quantitative analysis of H-bond distance was performed, shown in Fig 2. There are six H-bonds generate in complex from P-18S2 and PTX. It was worth to note that the combination between P-13S2 and PTX showed more H-bonds interaction from Bases of G1, G5, G9 and G13 and ligand, increased the stability of complex to a great extent (Fig 2). Therefore, P-18S2 showed a great activity to the obstruction of the binding between Sodium channel protein and PTX.

**Figure 2.**
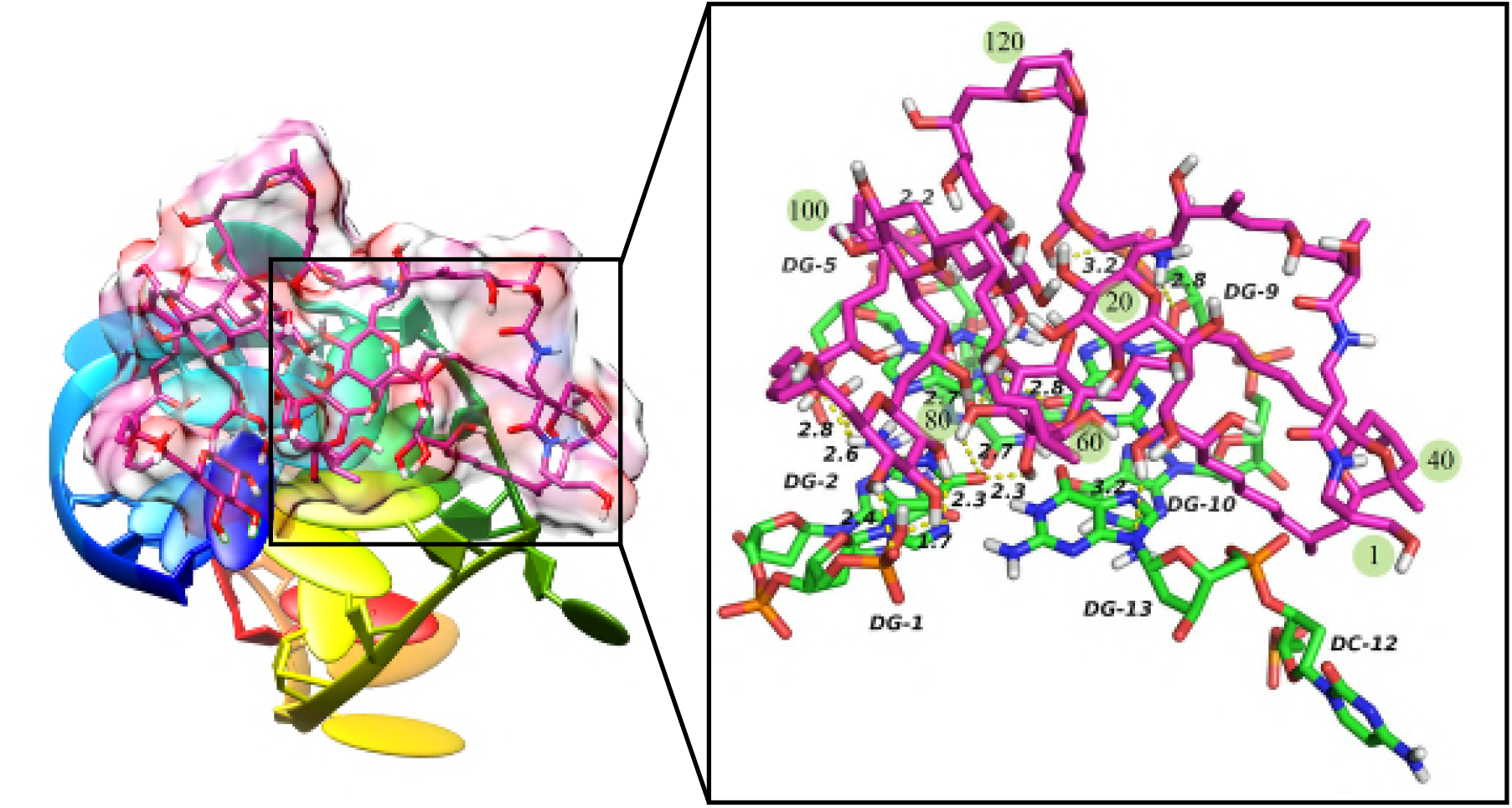
Three dimensional view of the interaction between P-18S2 and PTX.

### Analysis of molecular dynamics simulations

A common way to analyze the structural stability of a biomacromolecular in MD simulation is to monitor the root-mean-square deviation (RMSD) from the initial structure along the simulation. The RMSD of the complexes in this study are shown in Fig 3. It is clear from that plots that after 2ns the complex of P-18S2-PTX reached an equilibrium state with fluctuation of 2 Å. Furthermore, in 30 ns of simulation the structures were inconsiderably distinct from the initial structures that were employed as the starting point of the simulations, shown in Figs 4 and 5, indicating that the complexes were stable. The conformation of complex with different time from 0ns, 10ns, 20ns and 30ns were similar, without isolation from each other. The mass distance of G-quadruplex structures and PTX had also demonstrated it, shown in Fig 6. It is clear that there is no obvious fluctuation in the plots of mass distance between DNA and PTX versus simulation time. In addition, the binding energy (Fig 7) of complexes P-18S2-PTX kept equilibrated, indicated that there is no separation between the DNA and PTX, kept combined steadily. If there is dissociation, the energy will fluctuate severely. The absence of fluctuation showed the strong non-bond interaction between DNA and PTX, which made the complexes stable. Consequently, the high affinity between PTX and DNA P-18S2 blocked the binding between Sodium channel protein and PTX.

**Figure 3.**
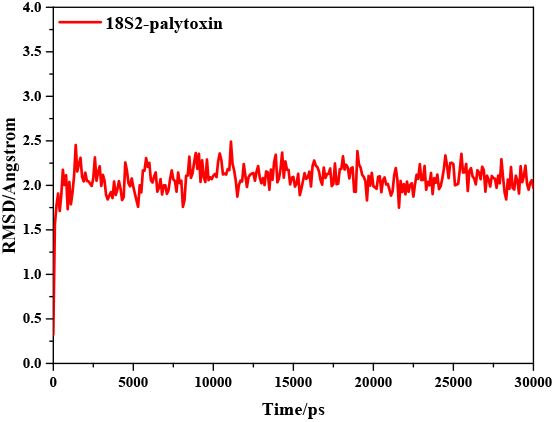
RMSD of CA atoms versus simulation time.

**Figure 4.**
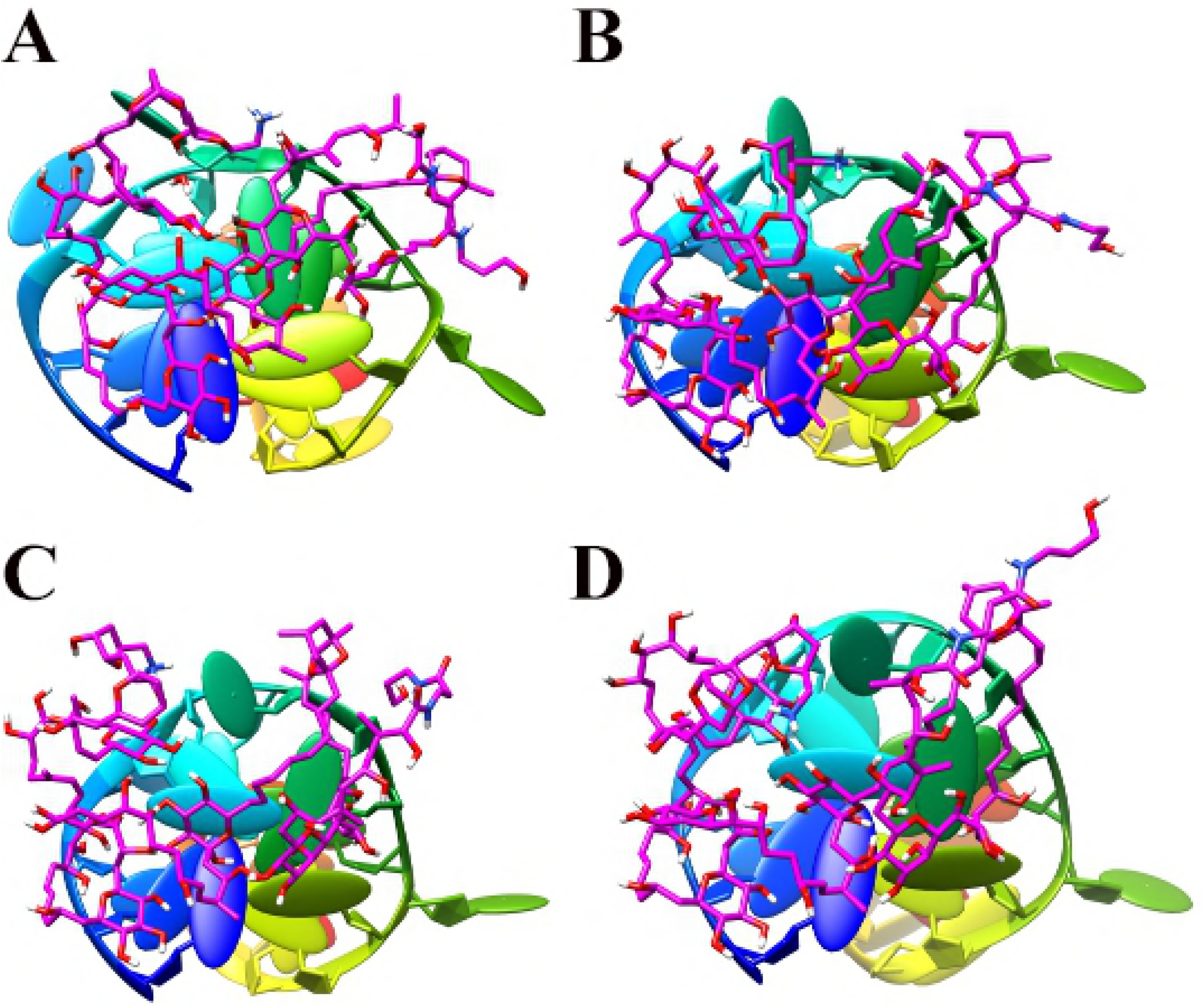
Top-view of the conformations of complex P-18S2-PTX versus MD simulation time A 0ns, B 10ns, C 20ns, D 30ns.

**Figure 5.**
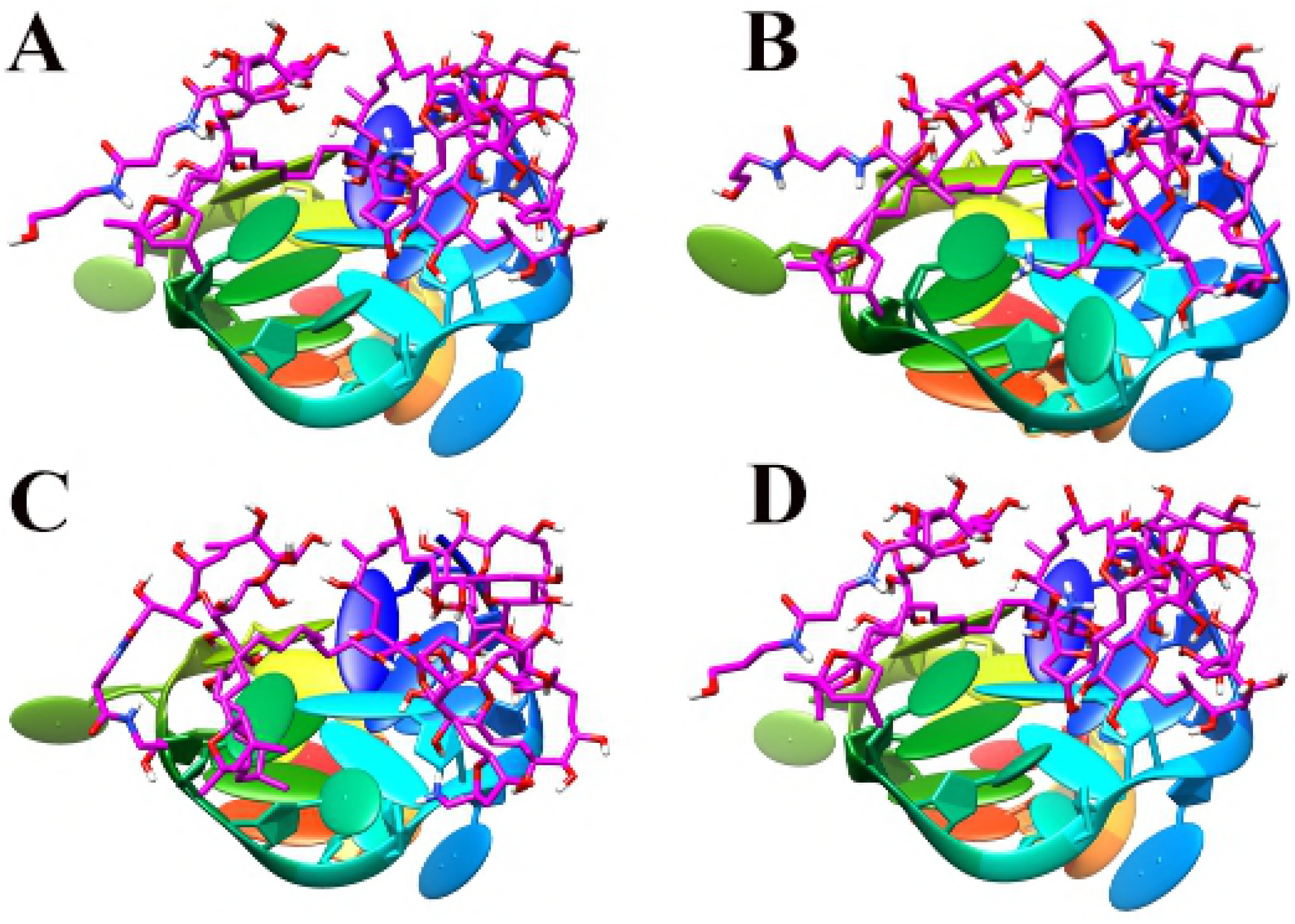
Oblique-view of the conformations of complex P-18S2-PTX versus MD simulation time A 0ns, B 10ns, C 20ns, D 30ns.

**Figure 6.**
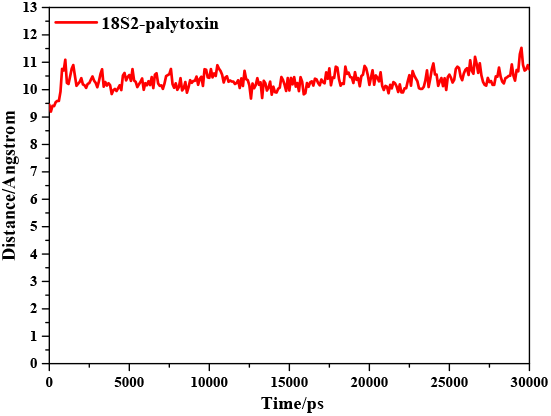
Mass distance between P-18S2 and PTX versus simulation time.

**Figure 7.**
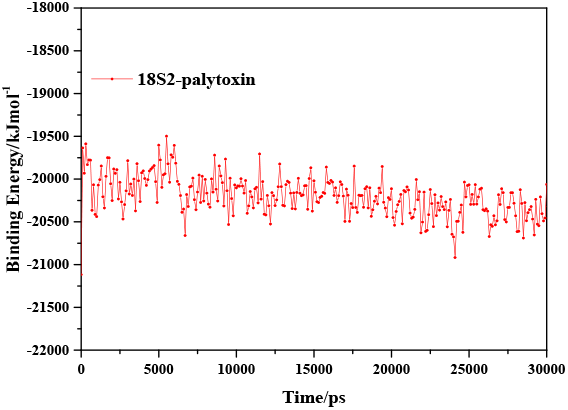
Binding energy of complexes versus simulation time.

The stability of G-quadruplex structures were analyzed as well, shown in S1 Fig. As we can see, the G-quadruplex structures of DNA P-18S2 show little fluctuation in the simulation, indicated that DNA P-18S2 with G-quadruplex structure has a strong intramolecular interaction, lead to the stability of G-quadruplex structure. As a result, the root mean square fluctuation (RMSF) (Fig 8) of the key bases of G-quadruplex structure, such as G1, G2, G5, G6, G9, G10, G13 and G14 in P-18S2, showed a small value less than 2Å. So it can be concluded that DNA with G-quadruplex structure keep steady in the progress of MD simulation, led to the stable combination between PTX and DNA. After quantitative analysis, it is found that the RMSF of the key bases of G-quadruplex structure (G1, G2, G5, G6, G9, G10, G13 and G14 with a value of 1.03, 0.91, 0.74, 0.68, 1.07, 1.21, 1.05 and 1.04 Å respectively) from P-18S2 is very low. That is why P-18S2 shows high activity for blocking the binding of PTX to Sodium channel protein.

**Figure 8.**
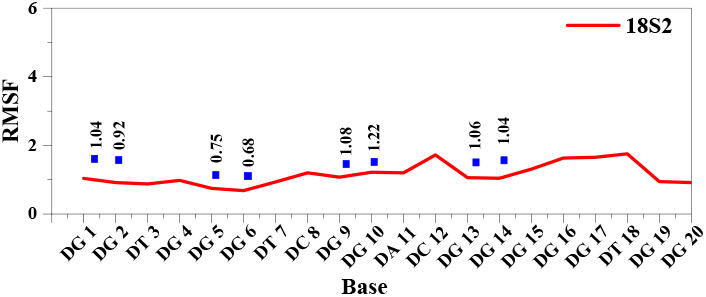
RMSF of residues of complex versus simulation time.

In order to give a new insight into the binding of DNA with PTX during the MD simulation, the hydrogen bonds were monitored, shown in Fig 9. The plots of the hydrogen bonds between DNA and PTX indicated that the DNA combined with PTX stably by the strong hydrogen bonds interaction. In the process of dynamic simulation, the hydrogen bonds of the complexes always exist. Some hydrogen bonds exist only for a period of time, named dynamic hydrogen bonds, while others, called static hydrogen bond, persisted through the MD simulations. The analysis of the dynamic behavior of hydrogen bonds was performed to explore the blocking mechanism of DNA for the combination of PTX and sodium channel protein. The three-dimensional structures of the interaction of DNA P-18S2 with PTX system was visualized in Fig 10. It is found that there are both static hydrogen bonds and dynamic hydrogen bonds between DNA P-18S2 and PTX during the dynamic simulation. As shown in Fig 10, there are three static hydrogen bonds and eighteen dynamic hydrogen bonds generated between P-18S2 and PTX, made the binding of P-18S2 and PTX more stable and more active to block of the binding of PTX to Sodium channel protein.

**Figure 9.**
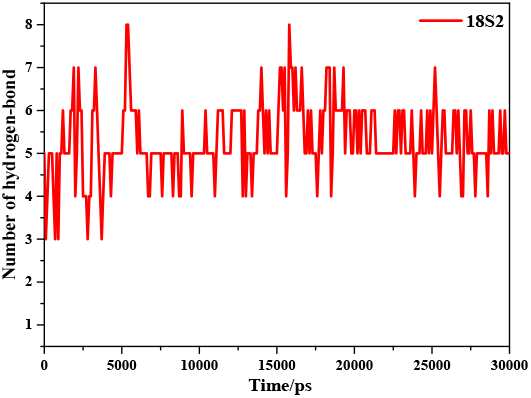
Hydrogen bonds between P-13S2, P-18S2 and PTX versus simulation time.

**Figure 10.**
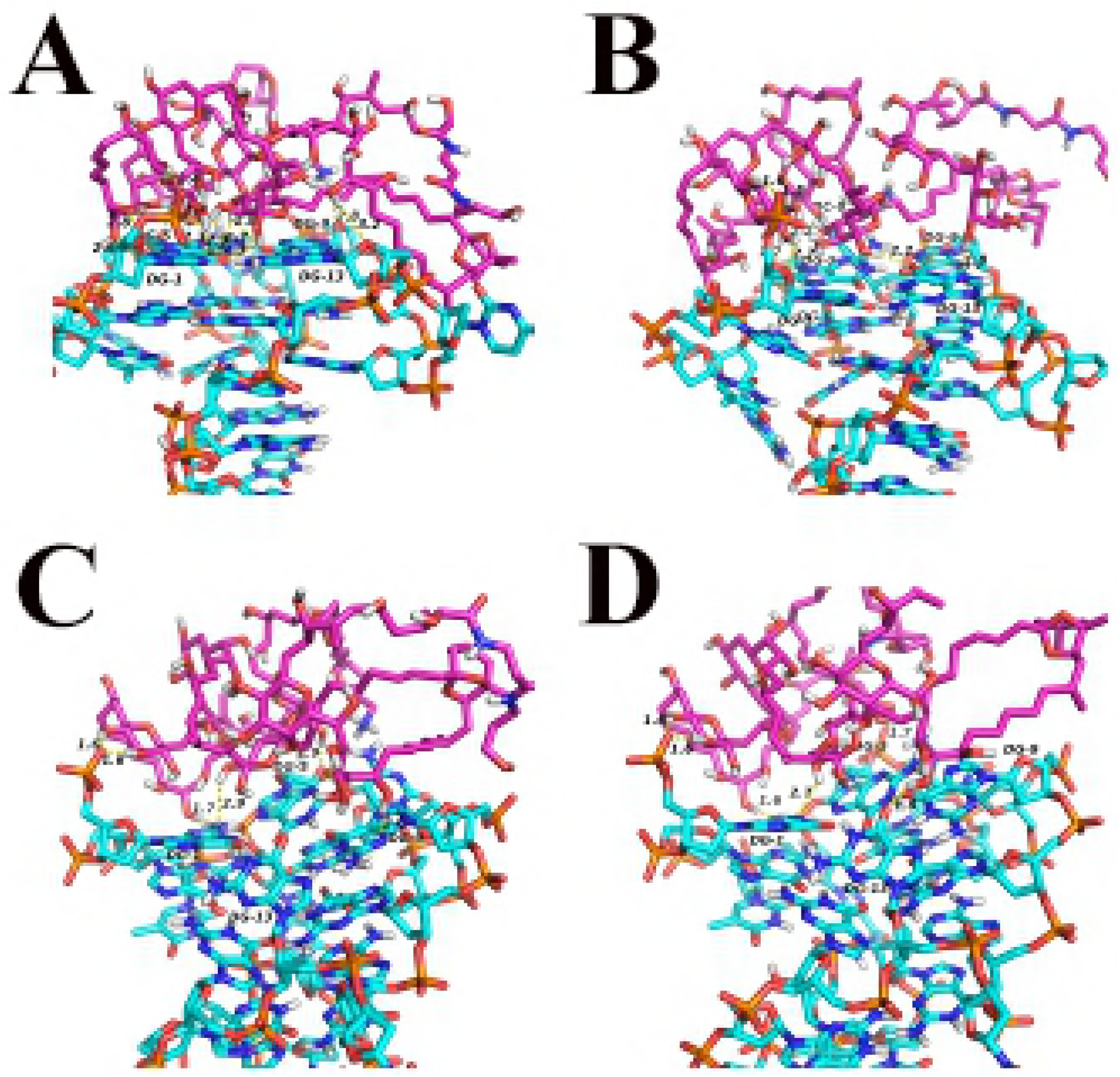
3D-view of hydrogen bonds interaction between P-18S2 and PTX versus simulation time A 0ns, B 10ns, C 20ns, D 30ns.

Strong hydrogen bonds interaction promoted the stability of the complexes. The trajectory diagram of radius of gyration also proved this point. The radius of gyration of biomacromolecular is a sign of stability in the process of dynamic simulation. As we can be seen from trajectory diagram of radius of gyration from the complexes P-18S2-PTX (Fig 11), radius of gyration is stable after 5 ns and the fluctuation value is negligible. Therefore, the radius of gyration of complex further validated the stable DNA with G-quadruplex structure and combination between DNA and PTX.

**Figure 11.**
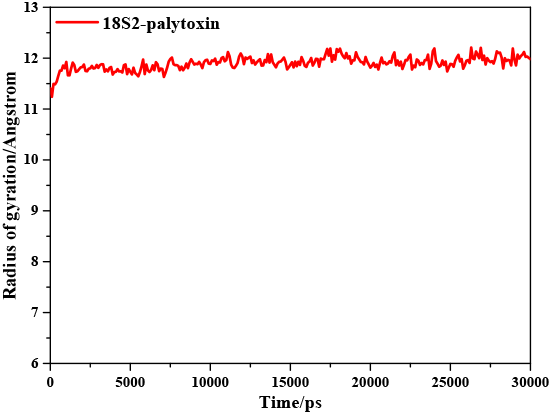
Radius of gyration of complex versus simulation time.

### Affinity of between truncation of P-18S2 and PTX

The BLI results showed that PTX interacted with truncated successively truncated sequence P18-S3,P18-S4, P18-S5, P18-S6, P18-S7, P18-S8 with a Kd (nM) of 3.14, 2.13, 0.87, 0.81, 2.93, 2.62 respectively, and P18-S2 with a Kd of 1.09 nM, shown in S2 Table and S2 Fig. Those data very close to the previous result(0.93 nM)^[10]^, and no phenomenon were observed in the corresponding blank or negative controls, which demonstrated that there is no significant difference between truncated P-18S2 and primary P-18S2 to bind PTX. In addition, this result further verified the reliability of computer simulation and defined the most core site of P-18S2 and PTX.

## Methods

### G-quadruplex model generation

The structure prediction of P-18S2 was performed with *QGRS(quadruplex forming G-rich sequences, http://bioinformatics.ramapo.edu/QGRS/index.php*)Minimum G-Group size was settled as 2. There are 56 possible structures for P-18S2 in forming G-quadruplex structure, the top 10 results of QGRS prediction were showed in S3 Table, with a maximum G-Score of 21. In a word, P-18S2 was great possibility in formation of G-quadruplex structure.

Then, the search in the Nucleic Acid Database (NDB)^[20]^for quadruplex DNA structures returned 193 entries. Unfortunately this list did not include all G-quadruplex containing structures. The Protein Data Bank (PDB)^[21]^ was searched as well for entries containing the words “quadruplex” and the list was filtered by visual inspection. The sequences extracted for each chain in the corresponding PDB files were aligned to P-18S2. The 2LXQ^[22]^ was found that it was exhibiting a corresponding intrastrand G-quadruplexes and similar size to P-18S2. In addition, the arrangement of guanine in 2LXQ was similar to P-18S2 to a certain extent. Based on the atomic models of 2LXQ, the 3 D model of P-18S2 with G-quadruplex structure was generated by Discovery Studio 2.5 Client (*http://accelrys.com/products/discovery-studio*) with the nucleic acids substitution, insertion and deletion.

Care was taken to make the coordination geometry most favorable. Conformations of a few nucleic acids were therefore adjusted manually within well allowed ranges. Then, the optimization of model was performed at the high performance computing facility with the YASARA package^[23, 24]^, using the Amber 14 force field^[25]^, and the water model was TIP3P. The temperature coupling of the model system was ascertained by Berendsen thermostat method, while the manometer method was used for pressure coupling^[26]^. Besides, the starting structure was immersed in a periodic rectangular simulation cubic cell of water. For the operation of optimization in the simulated water condition, the backbone was firstly fixed and the side chain was optimized 5,000 steps, and then, the whole structure was majorized 5,000 steps. Finally, the optimized 3D-struture of P-18S2 with G-quadruplex structure was generated, shown in Fig 12. After optimization, quantitative analysis was performed to the G-quadruplex structure of P-18S2. The results showed that, due to strong hydrogen-bonds interaction among G1, G5, G9, G13 and G2, G6, G10, G14 in P-18S2 (S3 Fig), formed stable G-quadruplex structure.

**Figure 12.**
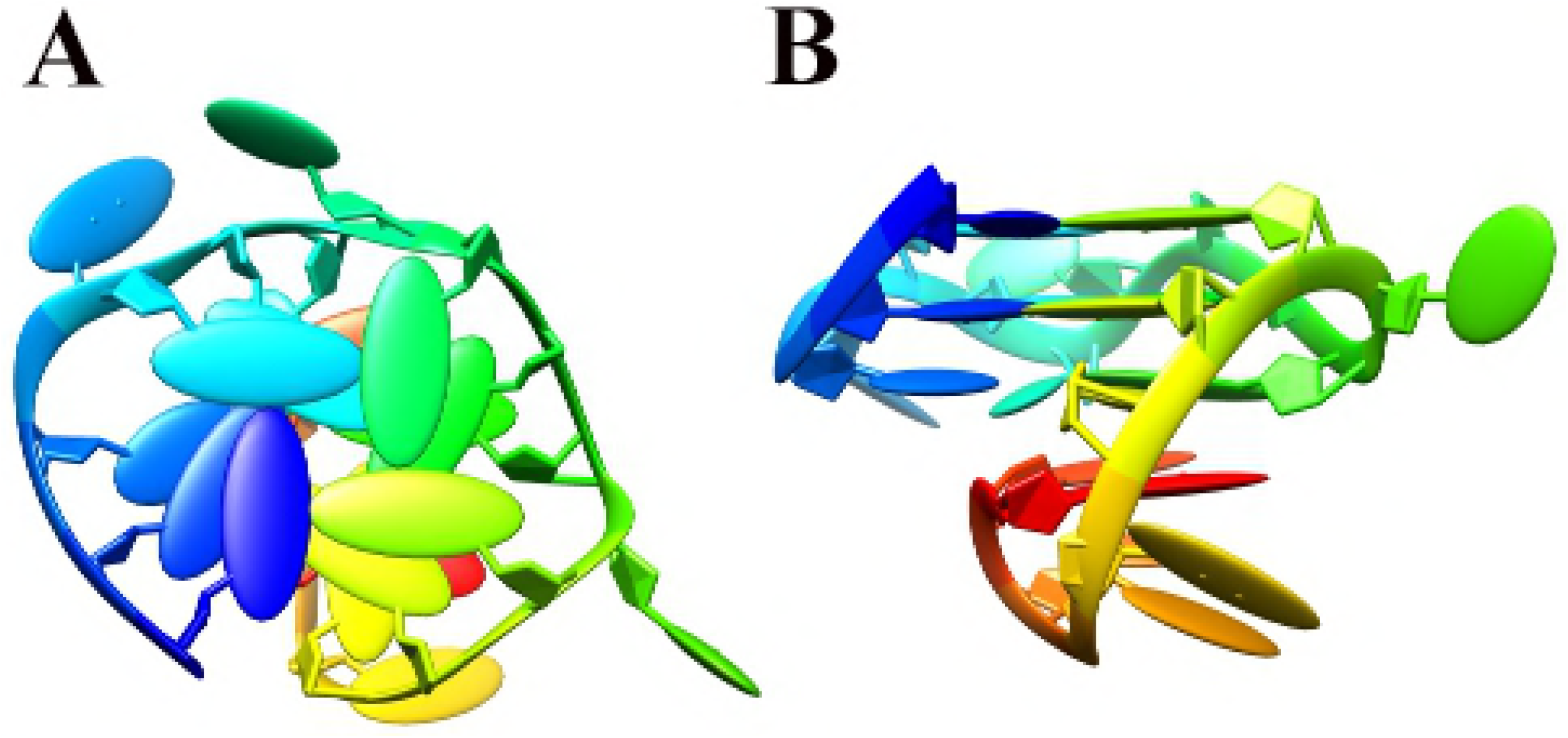
The 3D model of P-18S2 with G-quadruplex structure, top view at left and side view at right.

### Docking

Firstly, PTX was obtained from PubChem Compound database with accession no. 45027797, shown in S4 Fig. The PTX molecule was optimized using MM2 method. Due to large size and high molecular weight, PTX showed high flexibility and lots of possible conformations. So, it is unreasonable to perform the docking using PTX from the database directly. The solution to these problems is to first obtain the equilibrium structure of PTX in the solvent and then use it for molecular docking. Only in this way can we be able to investigate the interaction between PTX and DNA P-18S2 scientifically. In this work, 30 ns dynamics simulation is performed to PTX in KCl aqueous solution using GAFF force field^[27]^, then repeat three times. It was found that in the process of dynamic simulation the PTX tended to constriction in aqueous solution and reach an equilibrium state, shown in Fig 13. The 3D structures of PTX after equilibrium were output and optimized. Subsequently, the total energy was calculated, and the structure with lowest energy was selected to use for the subsequent simulations. Then, the number of rotatable bonds of ligands was set up in AutodockTools^[28]^. Finally, PTX was output and saved as the ligand.

**Figure 13.**
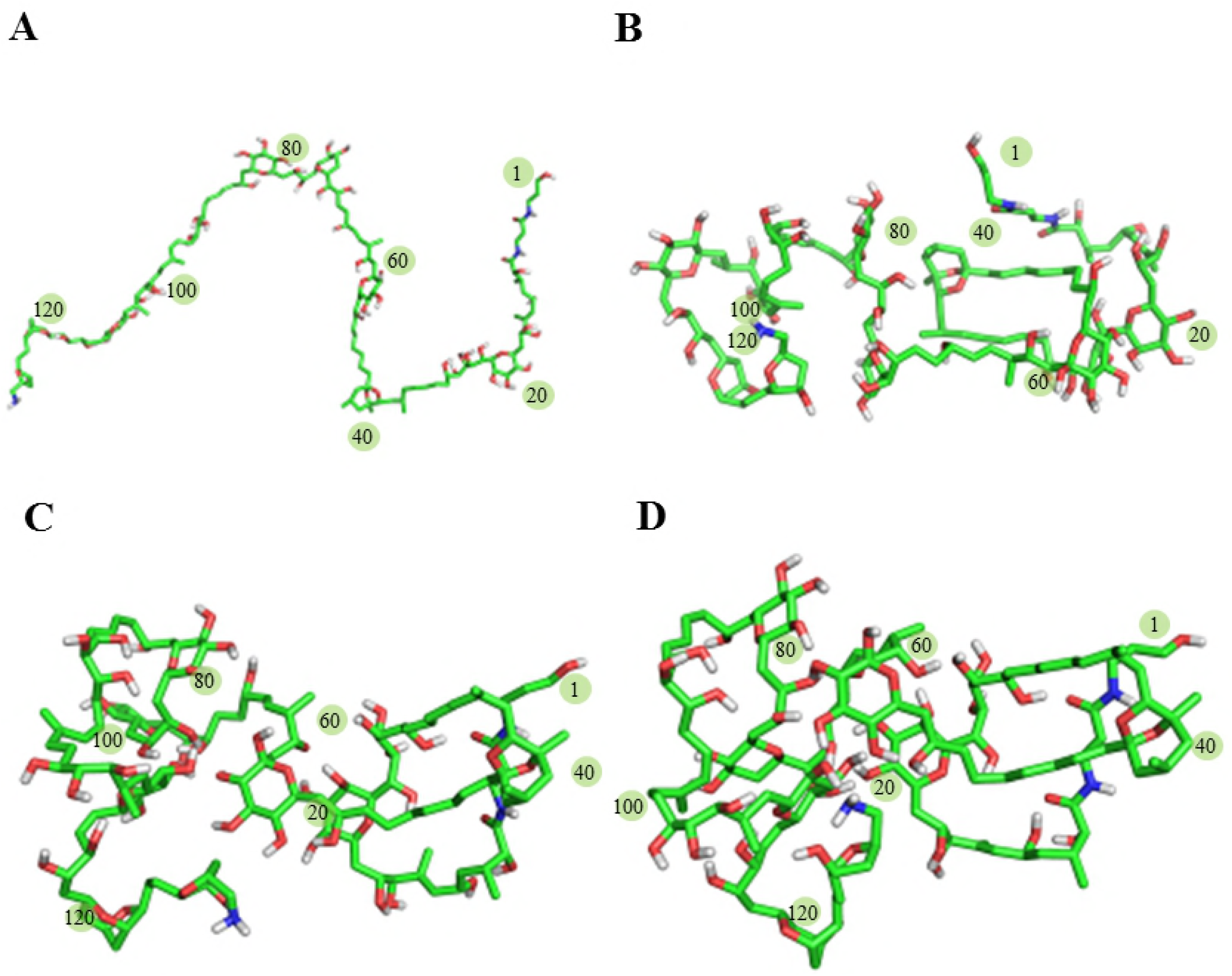
Conformation change of the PTX in the process of dynamic simulation :(A) 0ns, (B) 10ns, (C) 20ns, (D) 30ns.

For DNA-ligands docking, 3D structure of P-18S2 was obtained from the modelling and initialized as receptor molecules with AutodockTools. Subsequently, the receptor was endowed with AD atomic type, and hydrogen and charge were added, followed by the mergence of nonpolar hydrogen. The binding sites of P-18S2 recognizing ligand was obtained based on crystal structures of DNA-ligands complex with G-quadruplex structure^[29–33]^, which proved that the binding site of DNA with G-quadruplex structure were in the groove at the top and the bottom. In addition, the experiment suggested that the ligand could induce the formation of G-quadruplex structure. Therefore, as demonstrated in our previous workt^[17]^, the binding site of P-18S2 with G-quadruplex structure was at the top (S5 Fig).

Finally, the molecular docking analysis was carried out using Hex 8.0^[34]^. Rotation of receptor and ligand was allowed in 180 degree, without the specified binding group. The shape and electrostatic interaction were also considered. The maximum number of docking was revised as 200 and cluster analysis were made, while the default parameter settings were retained for others. For each of the docking cases, the lowest energy conformation, according to the Hex scoring function, was selected as the binding mode. The output from Hex was rendered with Chimera^[35]^ and PyMol program^[36]^.

### Molecular dynamics simulations

The Molecular dynamics (MD) simulations carried out were based on the complex which obtained from docking in Hex program. All simulations were performed using the molecular dynamics program YASARA and the amber 14 force field^[37]^. The complex using in the simulation came from molecular docking with hydrogen generated by the YASARA program. All simulations were carried out with an integration step of 1 fs and coordinates of the simulation model were recorded per 1ps. The starting structures were immersed in a periodic rectangular simulation cubic cell of KCl aqueous solution. The box dimensions were chosen to provide at least a 10 Å buffers of solvent molecules around the solute.

The fully solvated systems were then subjected to 5000 steps steepest descent minimization runs to remove clashes between atoms. An 80 ps position restrained MD simulation was performed for each system at constant pressure (1 atm) and temperature (300K). The temperature and pressure were kept constant during the simulations. Temperature coupling was done using the Berendsen thermostat with a temperature coupling constant of 0.1 ps, while the manometer method was used for pressure coupling, with a reference pressure of 1 atm. A particle mesh Ewald scheme^[38, 39]^ was used to calculate the long range electrostatic interactions with a 10 Å cutoff for the real space. A cutoff of 14 Å was used for the van der Waals interactions (Lennard-Jones terms).Translation and rotation corrections were enabled during MD simulations to ensure that structures in trajectory were well superimposed, which is convenient for the structure analysis. The chemical bond lengths involving hydrogen atoms were fixed with SHAKE algorithm^[40]^.

By the time of 1 ns, the simulation system reached an equilibrium state; thus, the system was subjected to conventional MD (CMD) simulation for 30 ns. All calculations were performed on the MolDesigner molecular simulation platform.

### Determination of affinity by BLI

We surprised to found that P18-S2 can be further truncated and optimized by simulating and docking. Therefore, the affinity of the successively truncated sequence (P18-S3~P18-S8) and P18-S2 for binding to PTX were determined respectively. The super streptavidin-coated (SSA) biosensor was used for the immobilization of the 7 sequences onto the BLI aptasensor. Those aptamers were analyzed for association time over 2 min with PTX(5 uM) and dissociation time over 2 min, along with a blank sample containing only binding buffer for reference and a random sequence was used as a negative control.

## Acknowledgements

We are sincerely grateful for Chengdu FenDi Technology on technical guidance in computer simulation.

## Supporting information

**S1 Fig. Oblique view of the comformations of G4 struture in DNA P-18S2 during MD simulationA 0ns, B 10ns, C 20ns, D 30ns.**

**S2 Fig. The red, green and blue lines respectivelyrepresent the interaction curve of aptamer18S2, 18S4,18S8 with PTX (5 uM).**

**S3 Fig. The G-quadruplex structure and intramolecular hydrogen bonds of P-18S2.**

**S4 Fig. Schematic representation of palytoxin.**

**S5 Fig. The binding site of P-18S2 (B) with G-quadruplex structure.**

**S6 Fig. The dynamic image of P-18S2 combined with PTX.**

**S1 Table. Docking results between DNA with G-quadruplex structure and palytoxin.**

^a^ The total calculated interaction energy;^b^ The original Hex steric complementarity score;^c^ The electrostatic energy;^d^ Root-mean-square deviation of the top-scoring docking orientation.

**S2 Table. The *K_d_* values of aptamers combined with PTX.**

**S3 Table. The results of P-18S2 from QGRS prediction.**

**S1 File. The README file.** This file includes all software and publicly available repositories along with our brief description.

**S2 File.** This file include input coordinates, topologies and parameter files.

